# Weak selection of phenotypic memory in periodic environments

**DOI:** 10.64898/2026.10.08.757665

**Authors:** Stephen R. Proulx, Henrique Teotónio

## Abstract

Offspring can inherit a physiological or molecular state from their parents, and the carry-over in phenotypes across generations that results is often called epigenetic or phenotypic “memory”. Most theory on the evolution of such memory asks how the rate of switching between pre-existing phenotypic variants evolves in a fluctuating environment. Here we ask instead how the variants themselves originate. We model an asexual population in a periodically alternating environment in which a genotype specifies two inherited states, the phenotypic value each produces and the probability with which each is retained from parent to offspring. Starting from an ancestor that produces a single phenotype, mutations creating small differences between inherited states have no first-order selective advantage. Their spread depends on second-order selection and requires strong selection against phenotype–environment mismatch. A maternal effect of the same small size, conditioned on the maternal environment, is under first-order selection in the same model. Once phenotypes have diverged, genotypes whose retention probabilities match the duration of the environmental states are strongly favored, and the evolved strategy persists under conditions in which it could not originate. Memory is a consequence of individual selection and its origin probably follows earlier selection for cue-dependent maternal effects.

## Introduction

Environments change, and a phenotype well suited to one set of conditions is often poorly suited to the conditions that the next generation faces. The resulting mismatch between the phenotypic values present in a population and the phenotypic value favored in the current environment imposes a fitness load that creates an opportunity for adaptation (Proulx and Phillips 2005). Populations can respond to such fluctuations in several ways, all of which alter how the phenotype of an individual is determined.

An individual’s phenotype depends on its genotype, on the environment experienced during development, through developmental plasticity (Via and Lande 1985; Moran 1992; Gavrilets and Scheiner 1993; Scheiner 1993), and on the inherited state received from its parents, typically its mother, in the form of a physiological condition or a molecular mark, through a maternal effect (Mousseau and Fox 1998; Badyaev and Uller 2009; Wolf and Wade 2009). When these one-generation effects occur generation after generation along a lineage, they generate a carry-over in terms of phenotype frequencies across generations, and the phenotype of an individual comes to resemble that of its parent, grandparent, and earlier ancestors beyond what a shared genotype and a shared environment would produce. This carry-over is what is usually meant by phenotypic “memory”, and when the inherited state is a chromatin modification or a small RNA, by epigenetic memory (Jablonka et al. 1995; Bonduriansky and Day 2009; Jablonka and Raz 2009; D’Urso and Brickner 2014; Jablonka 2017). Explaining how phenotypic memory evolves involves three separate problems, how a parent responds to an environmental cue, whether the physiological or molecular state that results is transmitted to the offspring as an inherited state, and whether the phenotype it produces in the offspring increases fitness (Uller 2008; Wolf and Wade 2009). The problems are separate even when one process contributes to more than one of them. Maternal provisioning that responds to the mother’s environment, for example, is a plastic response from the mother’s perspective and a maternal effect from the offspring’s perspective.

Interest in phenotypic memory has grown with reports that inherited states produced in response to the environment can persist for several generations (Heard and Martienssen 2014; Quadrana and Colot 2016; Fitz-James and Cavalli 2022), and with the idea that maternal effects prepare offspring for the conditions their mothers experienced (Gluckman et al. 2005; Marshall and Uller 2007; Storm and Lima 2010; Bell and Hellmann 2019). Meta-analyses of experiments in which offspring were raised in the same or in a different environment from their mothers find, however, that the benefit of a matched environment is small (Uller et al. 2013; Yin et al. 2019; Sánchez-Tójar et al. 2020). Most documented maternal effects, moreover, last a single generation (Galloway and Etterson 2007; Salinas and Munch 2012; Wibowo et al. 2016; Jordan et al. 2019; Perez and Lehner 2019), although in cases where inherited states do persist for longer there is heritable variation in the number of generations over which they persist (Frézal et al. 2018; Frejacques et al. 2026).

Experimental evolution offers a way to separate the three problems mentioned above, as our experiments with the nematode *Caenorhabditis elegans* illustrate. In oxygen environments that alternated between generations, populations adapted through a maternal phenotype, the provisioning of embryos with glycogen, that responds to the mother’s own environment. This is a plastic response of the mother and a maternal effect on the survival of her offspring in anoxia, and it is transmitted for a single generation (Dey et al. 2016). In oxygen environments that were uncorrelated between generations, the same kind of maternal effect evolved only when a feature of the mother’s environment other than oxygen reliably predicted the oxygen environment of the offspring, and populations without such a predictor became maladapted (Teotónio and Proulx 2026). In addition, over 60 generations of fluctuating oxygen, the dynamics of population growth were better described by an inherited state that responds to the environment and is retained across generations, incompletely and asymmetrically, than by developmental plasticity and maternal effects alone, so that transmission beyond one generation is present in these populations (Proulx et al. 2019). These experiments do not show, however, how the transmission of inherited states, state transmission for short, evolves from an ancestor that lacks it.

The strength of selection on a mutation that changes how the phenotype is determined depends on whether the change is conditioned on an environmental cue, be it external or internal to the organism. When a cue received by an individual or by its mother reliably predicts the environment that individual will experience, developmental plasticity and cue-dependent maternal effects can evolve through first-order selection, that is, selection whose strength is proportional to the size of the phenotypic effect of a mutation (Moran 1992; Leimar and McNamara 2015; McNamara et al. 2016; Proulx and Teotónio 2017). A mutation of small effect that moves the phenotype in the direction indicated by the cue increases geometric mean fitness in proportion to its effect (de Jong 1995; Lande 2014; Chevin and Lande 2015; Kuijper and Hoyle 2015; Tufto 2015). Consistent with such first-order selection, plasticity evolves rapidly under experimental evolution (Sikkink et al. 2014; Leung et al. 2020; Mallard et al. 2020), although the ability of plasticity to remove the fitness load might be limited by the reliability of cues and by the costs of plastic responses (Getty 1996; DeWitt et al. 1998). Random production of alternative phenotypes, or diversifying bet-hedging, which can itself be a maternal effect (Crean and Marshall 2009; Joschinski and Bonte 2020), can also be favored when mismatch is costly (Cohen 1966; Bull 1987; Seger and Brockmann 1987; Simons and Johnston 1997; Zhang and Hill 2005; Simons 2009, 2011; Scheiner 2014), but in a population that has evolved to the optimum among single-phenotype strategies, a small amount of randomization has no first-order effect on fitness. Its effect is instead proportional to the square of the difference in phenotypic value it introduces, a case of second-order selection. Diversification is then favored only when selection against mismatch is strong (Bull 1987; Proulx and Teotónio 2017).

Existing theory on phenotypic memory has not addressed whether state transmission is under first-order selection, like plasticity and cue-dependent maternal effects, or under second-order selection, like bet-hedging. One reason for this gap in the literature is that prior models assume that the organism is already capable of producing functionally different phenotypes that are aligned with the environmental variability that is present. For instance, models of switching between two inherited states show that the switching rate that evolves reflects the rate of environmental change (Ishii et al. 1989; Lachmann and Jablonka 1996; Kussell and Leibler 2005; Herman et al. 2014; Kuijper and Johnstone 2016). Experiments with engineered yeast switchers show that the switching rate that does best depends on the rate of environmental change (Acar et al. 2008; Stajic et al. 2022). Both begin with distinct phenotypes that are already favored in different environments and ask how long an inherited state should persist. Quantitative-genetic models of maternal inheritance, in which the maternal phenotype contributes to the offspring phenotype, describe the same carry-over for a continuous phenotype (Falconer 1965; Kirkpatrick and Lande 1989; Lande and Kirkpatrick 1990; Wolf et al. 1998). When the maternal contribution is allowed to evolve it comes to reflect the autocorrelation of the environment (Kuijper and Hoyle 2015), but the variation that it transmits is present from the outset (in effect encoded in the *G* matrix of genetic variances and covariances of phenotypic values).

Offspring retain the inherited state of their parent with some probability, and asking how this transmission evolves from an ancestor that lacks it requires describing it as the probability that an offspring retains the inherited state of its parent regardless of the environment of either, which we call a retention probability. Retention probabilities that do not depend on the environment separate the fitness consequences of transmitting an inherited state from those of responding to a cue (Shea et al. 2011; Day and Bonduriansky 2011), but they do not imply that inherited states are uncorrelated with the environment. Once inherited states produce different phenotypic values, differential survival or reproduction changes the distribution of inherited states entering transmission, and transmission preserves the association that selection generates. Before the inherited states produce different phenotypic values, however, they cannot be differentially selected and carry no information about the environment.

This raises the specific question we address in this article: can a genotype that transmits an inherited state be favored when the inherited states do not yet produce different phenotypic values? To address it, we develop a theoretical model in which a genotype specifies two phenotypic values, one for each inherited state, and two retention probabilities, which cover the range from phenotypic randomization, in which the inherited state of the offspring is independent of that of the parent, to retention of the inherited state in every generation. Starting from an ancestor that produces a single phenotypic value, for which retention has no effect on fitness, we ask whether mutations that create small differences between the phenotypic values of the two inherited states can invade, compare them with a cue-dependent maternal effect of the same size in the same environment, and follow the joint evolution of phenotypic values and retention.

## Methods

### Modeling framework

We consider an asexual population with discrete, non-overlapping generations experiencing periodic environmental fluctuations. There is no density or frequency dependence, so the long-term growth rate of each genotype can be calculated independently. A genotype can invade when its long-term growth rate is greater than that of the resident genotype (Lande 2007; Sæther and Engen 2015).

Each individual carries one of two inherited states, each of which produces a phenotypic value, and offspring may retain the inherited state of their parent. Offspring can maintain an inherited state or respond developmentally to the material and signals they receive (Wolf and Wade 2009; D’Urso and Brickner 2014; Fitz-James and Cavalli 2022). Retaining an inherited state does not mean that an entire phenotype is transmitted unchanged. An inherited state originating in the parent can instead alter the developmental processes that produce the offspring phenotype, and it is, in this sense, part of the offspring’s developmental environment. Accordingly, a retention probability can reflect a maternal phenotype affecting what is deposited, an offspring phenotype affecting maintenance or response, or both. Our model of an asexual population summarizes their consequences without distinguishing which individual determines the retention probabilities, because with asexual reproduction parent and offspring have identical fitness interests and the assignment of the phenotype to one or the other does not change the selection analysis (Wolf and Wade 2001).

### Notation

We index genotypes by *g*. Each genotype is characterized by the two phenotypic values it can produce and by the transmission matrix *B*_*g*_ that specifies state transmission between parent and offspring. We collect these four quantities in the strategy vector ***θ***_*g*_ (Table 1).

**Table 1.** Notation.

| Symbol | Definition |
| --- | --- |
| $g$ | Genotype index. Resident and mutant genotypes are denoted by $r$ and $m$, respectively. |
| $i$ | Index for the two inherited states within a genotype, $i \in \{1, 2\}$. |
| $e$ | Index for the two environmental states, $e \in \{1, 2\}$. |
| $E_t$ | Environmental state experienced in generation $t$. |
| $k$ | Number of consecutive generations spent in each environmental state. A complete cycle lasts $2k$ generations. |
| $\rho$ | Lag-one autocorrelation of the periodically alternating environment, $\rho = (k - 2)/k$. |
| $z_{gi}$ | Phenotypic value expressed by genotype $g$ when it carries inherited state $i$. |
| $z_e^*$ | Optimal phenotype in environment $e$, with $z_1^* = -1$ and $z_2^* = 1$. |
| $\beta_{gi}$ | Probability that offspring of genotype $g$ retain inherited state $i$ of their parent (retention probability). |
| $\theta_g$ | Phenotype and transmission strategy of genotype $g$, $(z_{g1}, z_{g2}, \beta_{g1}, \beta_{g2})$. |
| $B_g$ | Transmission matrix of inherited states for genotype $g$. Rows identify the inherited state of the parent and columns that of the offspring. |
| $q_i$ | Stationary frequency of inherited state $i$ under transmission alone. |
| $h$ | Lag-one correlation between the inherited states of parent and offspring, $h = \beta_{g1} + \beta_{g2} - 1$, the phenotypic memory generated by $B_g$. |
| $w_e(z)$ | Fitness of phenotype $z$ in environment $e$. |
| $\eta^2$ | Strength of selection against mismatched phenotypes. |
| $W_{e,g}$ | Diagonal fitness matrix for genotype $g$ in environment $e$. |
| $\mathbf{n}_{g,t}$ | Row vector giving the abundance of genotype $g$ in each inherited state in generation $t$. |
| $M_g$ | Population projection matrix for genotype $g$ over one complete environmental cycle. |
| $\Lambda_g$ | Dominant eigenvalue of $M_g$, giving growth over one complete environmental cycle. |
| $\lambda_g$ | Per-generation log growth rate, $\lambda_g = 1/(2k) \log \Lambda_g$. |

Inherited states are indexed by *i* ∈ {1, 2}, and *z*_*gi*_ denotes the phenotypic value produced by genotype *g* in inherited state *i*. The labels are defined separately within each genotype, so inherited state 1 of genotype *a* need not correspond to inherited state 1 of genotype *b*. This notation allows us to compare coexisting genotypes or successive genotypes along a mutational sequence.

The phenotype and transmission strategy of genotype *g* is

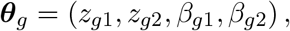

where *β*_*gi*_ is the probability that an offspring retains inherited state *i*, conditional on its parent carrying inherited state *i*. We refer to *β*_*g*1_ and *β*_*g*2_ as the retention probabilities, or transmission probabilities, of genotype *g*, and to the pair as its transmission strategy. The transmission matrix of genotype *g* is

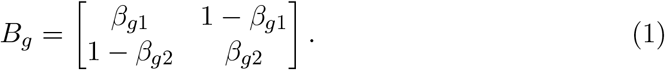

Rows of *B*_*g*_ correspond to the inherited state of the parent and columns to that of the offspring. Therefore,

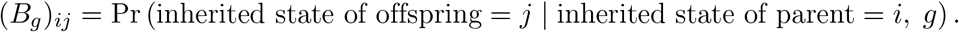

We use *e* ∈ {1, 2} to index the two environmental states and *E*_*t*_ to denote the environment in generation *t*. We consider a cyclic environmental process in which each environment remains constant for *k* generations before switching to the other environmental state. A complete cycle therefore lasts 2*k* generations.

The optimal phenotype in environment *e* is 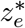, where we have arbitrarily set

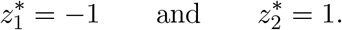

The fitness of phenotype *z* in environment *e* is

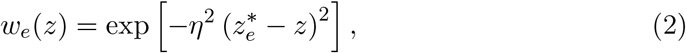

where *η*^2^ determines the strength of selection against mismatched phenotypes. In the usual Gaussian parameterization *η*^2^ = 1*/*(2*σ*^2^), where *σ* is the width of the fitness function.

We can express fitness for genotype *g* in matrix form as

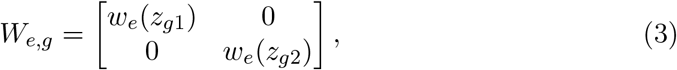

where the diagonal elements give fitness of genotype *g*’s phenotypes in environment

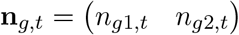

be a row vector giving the abundance of genotype *g* in each inherited state in generation *t*. Selection is followed by transmission of the inherited state, giving

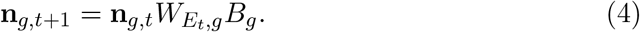

Without loss of generality, we define the beginning of an environmental cycle as the start of environment 1. The population projection matrix over one complete cycle is

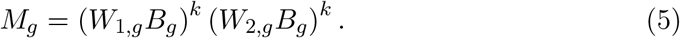

We denote the dominant eigenvalue of *M*_*g*_ by Λ_*g*_, the growth multiplier over one environmental cycle. Note that changing the order of the environments does not change the dominant eigenvalue. The equivalent per-generation log growth rate is

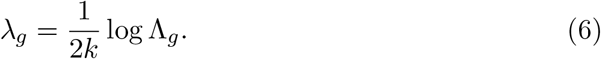

A mutant genotype *m* invades a resident genotype *r* when

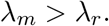

When only one genotype is under consideration, we suppress the genotype subscript.

### Transmission, stationary frequencies of inherited states and phenotypic memory

For a fixed genotype, equation 4 is a first-order Markov process, in which the inherited state of the parent carries all inherited information relevant to offspring development and carry-over over several generations follows from repeated one-generation transmission. A measured phenotype need not capture every inherited component that affects offspring development, and we return to this point in the Discussion.

We can summarize the inheritance process by the lag-one correlation *h* between the inherited states of parent and offspring (in the absence of selection). To characterize transmission separately from selection, we consider the stationary distribution of inherited states within a genotype in the absence of differential survival or reproduction. The stationary frequencies of the two inherited states are then

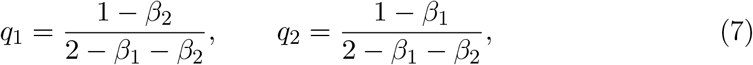

and the lag-one correlation is

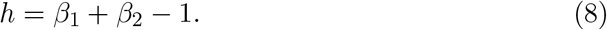

Note that now,

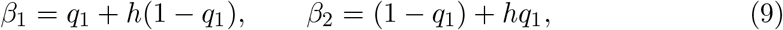

so that the transmission strategy can equivalently be described by the stationary frequency of an inherited state and its lag-one correlation, the two quantities that a phenotypic time series can estimate. The feasible range of *h* depends on the values of *β*_1_ and *β*_2_, and strongly negative correlation is possible only when the two inherited states have similar frequencies. The correlation between inherited states *l* generations apart is *h*^*l*^. Such carry-over is what we call phenotypic memory. It is a property of the transmission probabilities, not a phenotype expressed by any individual, and observed correlations can be altered by selection or by correlated environmental inputs.

The transmission matrix spans the range of inheritance patterns that a two-state system can display. When *β*_1_ = 1 − *β*_2_, the two rows of *B* are identical, the inherited state of the offspring is independent of that of the parent (*h* = 0), and the genotype produces the two inherited states at fixed frequencies, which is phenotypic randomization, or diversifying bet-hedging (Bull 1987; Proulx and Teotónio 2017). When both *β*_1_ and *β*_2_ are close to one, each inherited state is almost always retained, as a genetic locus would be with a low mutation rate. Intermediate values describe inherited states that are retained incompletely, and 1 − *β*_1_ and 1 − *β*_2_ can equally be read as switching or mutation rates (Ishii et al. 1989; Lachmann and Jablonka 1996; Kussell and Leibler 2005).

### The ancestral single-phenotype strategy

Because we want to understand how phenotypic memory can arise from an ancestor that lacks it, we consider an ancestral population that produces a single phenotype. A single phenotype that improves fitness in some of the experienced environments and results in a net increase in the geometric mean fitness of the genotype is a form of “conservative bet-hedging” (Seger and Brockmann 1987), and it can evolve by small steps, because successive mutations that increase geometric mean fitness each spread under selection. Selection of this kind is “first-order”, in that the selection coefficient changes by an amount proportional to the change in phenotype when the phenotypic change is small.

When both inherited states of a genotype produce the same phenotypic value,

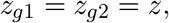

the entries of *B*_*g*_ have no effect on fitness because switching between the two inherited states does not change the expressed phenotype. The per-generation log growth rate is the logarithm of the geometric mean fitness in the two environments,

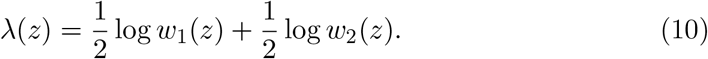

Substituting the fitness function from equation 2 gives

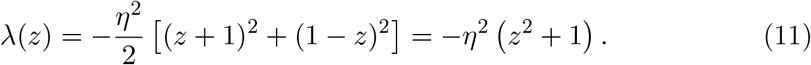

Differentiating equation 11 with respect to *z* gives the ancestral single-phenotype equilibrium

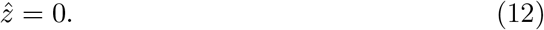

The log growth rate at this equilibrium is

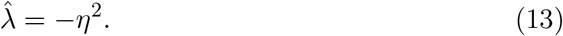

This calculation is what we use to define the single-phenotype ancestral strategy as the resident population for the initial invasion analyses.

### Invasion analysis

We can calculate mutant log growth, the invasion fitness of the mutant, from equations 5 and 6. To determine whether or not a mutation of small effect would spread we differentiate *λ*_*m*_ with respect to the changing parameter and evaluate the sign and magnitude of the derivative. When the first derivative exists and is zero, selection is weak, and we evaluate the second derivative to determine whether small changes in either direction are favored.

### Numerical simulations

We used deterministic simulations to examine how phenotypic values and retention probabilities change together. These simulations describe a sequence of changes through the four-dimensional strategy space

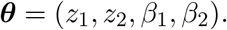

We approximate evolutionary change by moving up the fitness gradient such that each change is made in the direction that produces the largest local increase in *λ*.

For each environmental run length *k*, we constructed the cycle matrix *M*, calculated its dominant eigenvalue, and used equation 6 to obtain *λ* as a function of
the four components of ***θ***. We then calculated the gradient

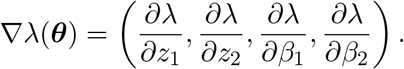

At each evolutionary step, we evaluated all four components of this gradient, identified the parameter with the largest absolute derivative, and changed that parameter. To constrain the phenotypic values to (−1, 1) and the retention probabilities to (0, 1), changes were made on logit scales adapted to the two ranges. For mutations in the *z* parameters we used the formula

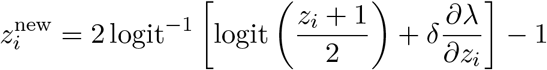

and for parameters on the 0-1 scale we used

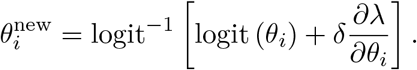

Here *δ* is an adjustable parameter controlling the mutation step size. All other components of ***θ*** remained unchanged during that step. We set *δ* = 1.

The simulation continued until the largest absolute component of the gradient was less than 10^−4^. The number of simulation steps does not correspond to a number of biological generations. Instead, the simulations describe the order in which the four components of the strategy change under repeated directional selection.

At each evolutionary step, we recorded the four strategy parameters, the pergeneration log growth rate, and the four components of the fitness gradient. We summarized the overall magnitude of the local gradient as

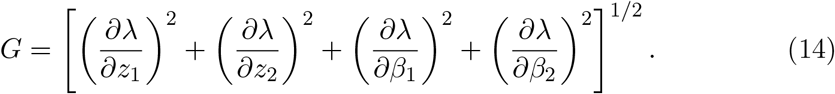

### Association between phenotype and environment

We quantified the association between inherited state and environment along each evolutionary trajectory using mutual information. For each strategy, we first found the periodic distribution of the two inherited states. We began with the normalized dominant left eigenvector of the cycle matrix and iterated the population recursion through one complete environmental cycle, normalizing the frequencies of the inherited states after each generation.

Let *f*_*i*|*e*_ denote the overall frequency of individuals in inherited state *i* during generations in which the environmental state is *e*. If *p*_*e*_ is the frequency of environmental state *e*, then

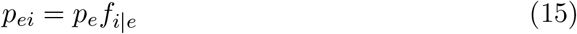

is the joint frequency of environmental state *e* and inherited state *i*. Equivalently, *p*_*ei*_ is the probability that an individual sampled from the population at a randomly sampled generation carries inherited state *i* while the environmental state is *e*.

Mutual information between environmental state and inherited state was calculated as

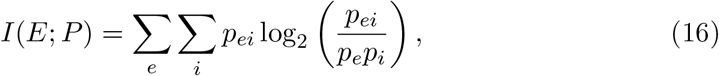

where *p*_*i*_ = ∑_*e*_ *p*_*ei*_ is the overall frequency of individuals in inherited state *i*.

The largest attainable association is less than the information-theoretic maximum of one bit because an inherited state cannot change until the generation after an environmental transition. We therefore scaled mutual information by the value obtained when phenotype matches the environment throughout each environmental run except for the first generation after each transition. The frequency of phenotype– environment mismatch under this strategy is 1*/k*, giving

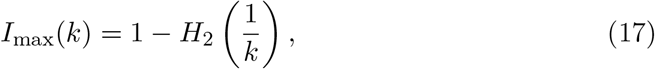

Where

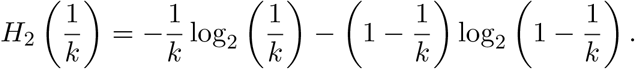

Relative information was calculated as

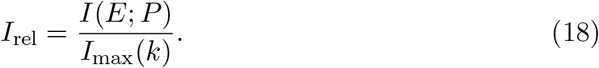

### Data accessibility

All symbolic and numerical calculations were carried out in *Mathematica* 14.0 (Wolfram Research, Inc. 2024). The notebooks that reproduce the analyses and the figures will be archived at Zenodo, currently available through github https://github.com/srproulx/PeriodicPhenotypicTransmission.

## Results

### Invasion of a mutant that randomizes the phenotype

We first considered mutations that introduce phenotypic variability without state transmission. Starting from the ancestral equilibrium at 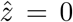, these mutants produce the ancestral phenotype, 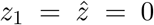, in one inherited state and an alternative phenotype, *z*_2_ = *ϵ*, in the other. We set *β*_1_ = *β* and *β*_2_ = 1 − *β*, making the inherited state of the offspring independent of that of the parent (*h* = 0). The alternative phenotype is produced with probability 1 − *β*, and values of *β* near one describe genotypes that produce it only rarely. Because the two environments and fitness functions are symmetric, choosing the alternative phenotype to be −*ϵ* would produce an equivalent analysis.

With *β*_1_ = *β* and *β*_2_ = 1 − *β*, the two rows of the transmission matrix in equation 1 are identical, and the frequencies of the two inherited states among offspring are *β* and 1 − *β*. The mutant’s multiplicative growth factor in environment *e* is therefore

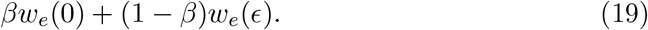

Because the two environments occur at equal frequencies, the mutant’s per-generation log growth rate reduces to

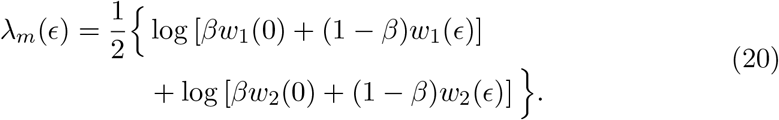

Note that this expression is independent of the environmental run length *k* because it depends only on the frequency, not the order, of the environmental states. Differentiating equation 20 with respect to *ϵ* and evaluating at *ϵ* = 0 gives

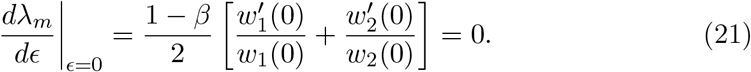

We have already found that 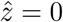 is the point at which the two log-fitness derivatives cancel. Thus, mutations that solely create phenotypic variability have no first-order fitness effect. We can evaluate their invasion fitness for slightly larger mutational effects by evaluating the second derivative,

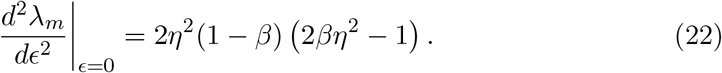

For 0 ≤ *β <* 1, the second derivative is positive when

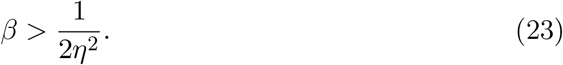

Because *β* ≤ 1, phenotypic variability cannot be favored unless *η*^2^ *>* 1*/*2. Put another way, if *η*^2^ ≤ 1*/*2 then phenotypic variability is not favored, regardless of the value of *β*. If, however, *η*^2^ *>* 1*/*2, then for high enough values of *β*, phenotypic variability is selected for. Genotypes that are most likely to produce the ancestral phenotypic value and only rarely produce the alternative variant are favored when *η*^2^ is above, but near, 1*/*2. For higher values of *η*^2^, phenotypic variability is favored even when both phenotypic variants are produced at higher frequencies. The threshold reproduces our earlier result for diversifying bet-hedging (Proulx and Teotónio 2017).

When selection is only slightly stronger than this threshold, invasion is limited to genotypes that produce the ancestral phenotype with high probability and the alternative phenotype only rarely. As selection against phenotypic mismatch becomes stronger, a broader range of values of *β* permits invasion of variable phenotype mutants (Figure 1).

**Figure 1.**
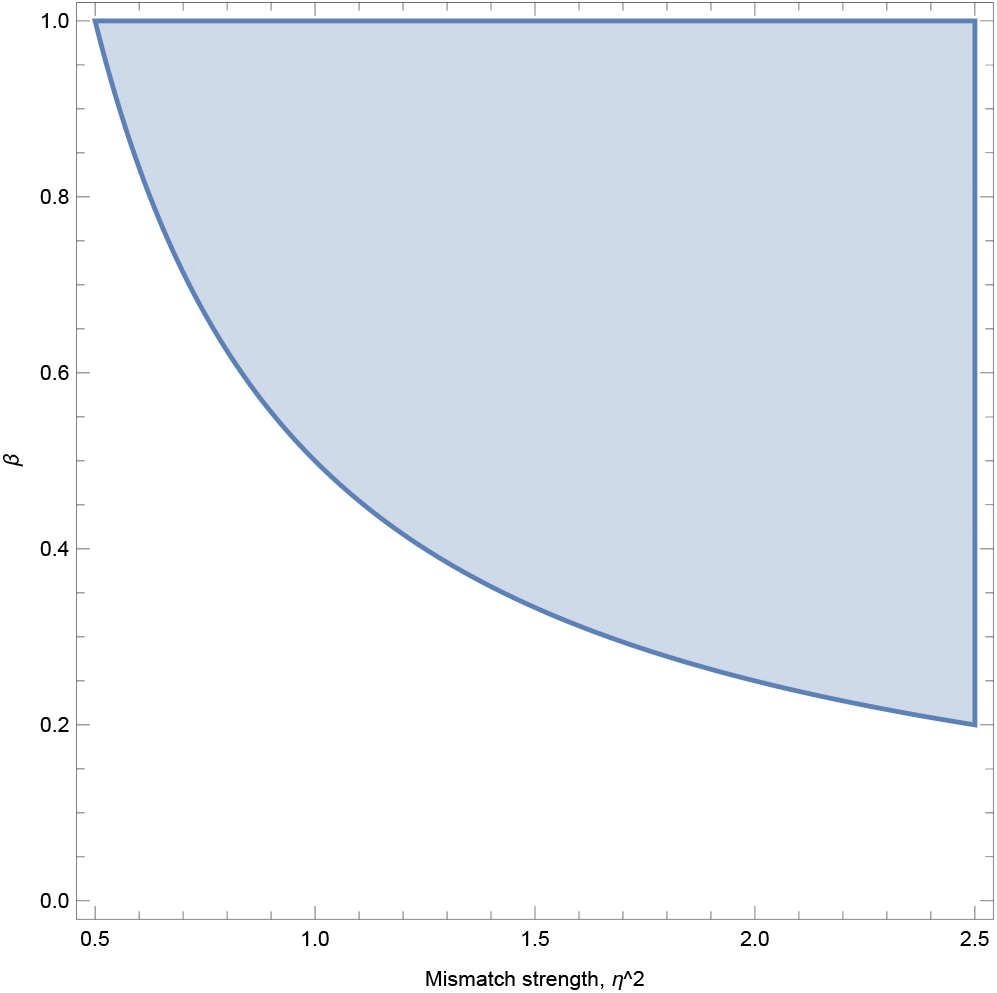
The region of parameter space in which phenotypic randomization can invade. Shading indicates combinations of the probability *β* of producing the ancestral inherited state and the strength of selection against phenotype–environment mismatch, *η*^2^, for which the second derivative of invasion fitness is positive. High values of *β* correspond to rare production of the alternative phenotype.

### Invasion of a mutant with state transmission

We next considered the invasion of a mutation that creates state transmission near the ancestral single-phenotype strategy, *z*_1_ = *z*_2_ = 0. For a mutant with *z*_1_ = 0, *z*_2_ = *ϵ* and general retention probabilities *β*_1_ and *β*_2_, we write the dominant eigenvalue of the cycle matrix *M*_*m*_ (equation 5) as a function of *ϵ*, Λ(*ϵ*), and derive its first and second derivatives at *ϵ* = 0 from the characteristic equation of this 2 × 2 matrix,

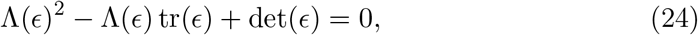

where tr(*ϵ*) and det(*ϵ*) are its trace and determinant. In terms of the per-generation log growth rate, 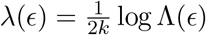, equation 24 reads

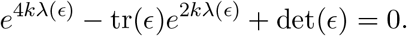

At *ϵ* = 0,

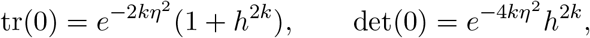

which gives *λ*(0) = − *η*^2^, the result obtained by direct calculation in equation 13 (See *Mathematica* notebook https://github.com/srproulx/PeriodicPhenotypicTransmission).

Implicit differentiation of the characteristic equation and solution for Λ′(0) gives

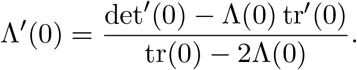

Both det′(0) and tr′(0) are zero, and therefore

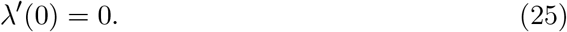

Regardless of the cycle length, the retention probabilities and the strength of selection, there is thus no first-order selection on state transmission at its origin. Note that at *z*_1_ = *z*_2_ the derivatives of *λ*_*m*_ with respect to *β*_1_ and *β*_2_ are also identically zero, because every transmission matrix produces the same expressed phenotype. Selection on state transmission is therefore at best second order. Differentiating the characteristic equation a second time, substituting tr′(0) = 0 and *λ*′(0) = 0, solving for *λ*″(0) and expressing the result in terms of *h, q*_1_ and *q*_2_ yields

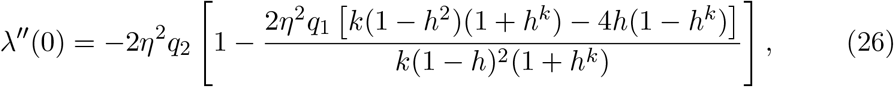

which reduces to equation 22 when *h* = 0 and *q*_1_ = *β*. The values of *β*_1_ and *β*_2_ for which *λ*″(0) is positive are those for which state transmission is weakly selected for.

### Comparison with a deterministic maternal effect

A deterministic maternal effect (DME) sets the offspring phenotype according to the maternal environment without error (Proulx and Teotónio 2017), whereas state transmission retains the inherited state of the parent with probabilities that do not depend on the environment.

To compare the two within the same environmental and fitness model, consider a deterministic maternal-effect mutant that produces

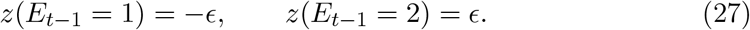

Positive *ϵ* moves offspring toward the optimum associated with the maternal environment. During each environmental run, maternal and offspring environments are the same for *k* − 1 generations and differ in the first generation following the environmental transition. The mutant’s per-generation log growth rate is therefore

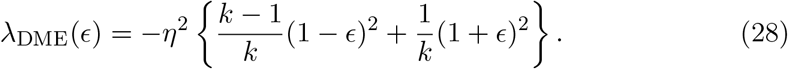

At *ϵ* = 0, this reduces to the ancestral log growth rate 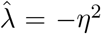. Differentiating gives

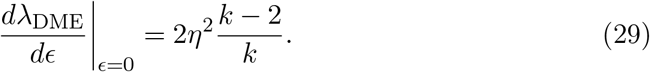

The factor (*k* − 2)*/k* is the lag-one autocorrelation of the two-state environment. We denote this environmental autocorrelation by *ρ* = (*k* − 2)*/k*. It is positive when *k >* 2, zero when *k* = 2, and negative when *k* = 1. When environmental runs last more than two generations, an arbitrarily small maternal effect is favored by first-order selection. When *k* = 1, the maternal environment perfectly predicts the opposite offspring environment, and selection favors a reversed maternal effect (*ϵ <* 0). When *k* = 2, the maternal environment contains no net one-generation predictive information, so the first derivative is zero.

The locally optimal deterministic maternal effect is obtained from equation 28:

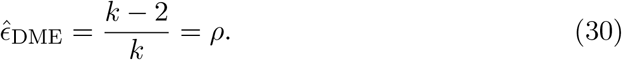

This gives the maximal log growth of

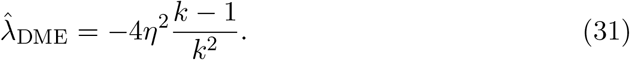

The optimum does not place offspring at the optimum of the maternal environment because doing so would impose a large cost in the transition generation. Instead, the phenotype is displaced in proportion to the predictive value of the maternal environment.

Figure 2 compares the geometric mean fitness, *e*^*λ*^, of the two kinds of mutant near the ancestral equilibrium as a function of the size of their phenotypic perturbation. The fitness of the deterministic maternal-effect mutant increases in proportion to *ϵ*, whereas that of the state-transmission mutant, which includes the second-order effects, increases only in proportion to *ϵ*^2^ and remains close to the ancestral value over the same range. The difference is not in what is transmitted but in whether the phenotypic effect of the mutation is conditioned on an informative cue at the moment it arises.

**Figure 2.**
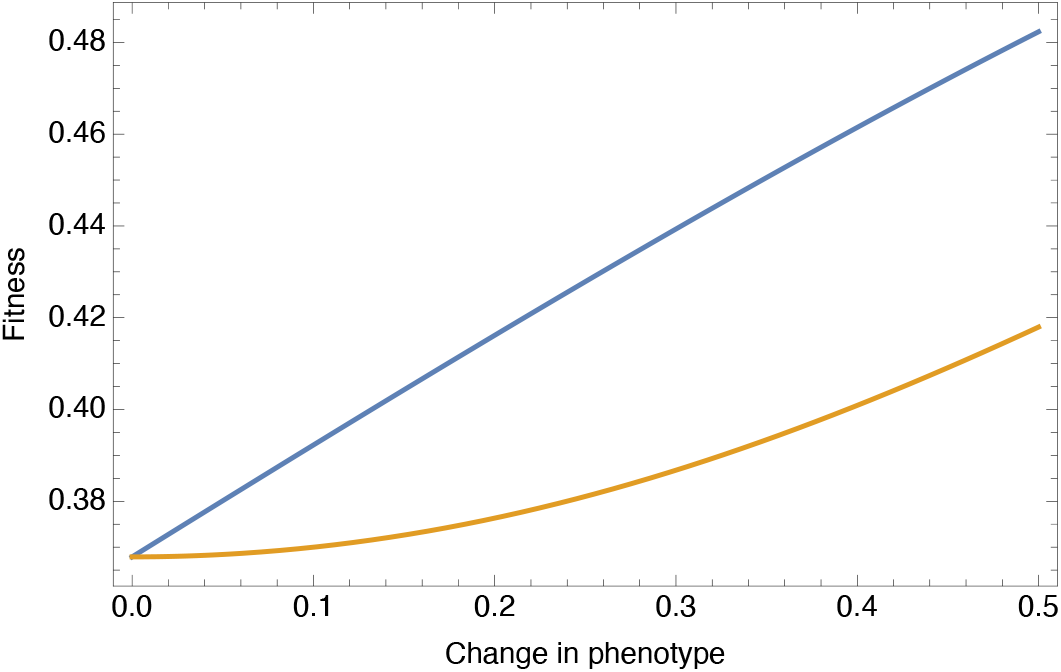
Geometric mean fitness of a deterministic maternal-effect mutant and of a state-transmission mutant near the ancestral single-phenotype strategy. Mutant geometric mean fitness, *e*^*λ*^, is shown as a function of the perturbation difference between the phenotypes produced for a deterministic maternal effect that conditions the offspring phenotype on the maternal environment (equation 28, upper blue curve) and for a state-transmission mutant with environment-independent retention probabilities. Both curves start at the ancestral fitness. The maternal effect is under first-order selection and its fitness rises linearly, whereas the fitness of the state-transmission mutant changes little until the difference in phenotypic value becomes larger. Parameter values: *k* = 6, *η*^2^ = 1, *β*_1_ = *β*_2_ = 5*/*6.

### Pre-existing transmission structure determines when phenotypic variability can invade

Even though state transmission does not generate first-order selection on a new phenotypic difference, second-order selection can be positive when the transmission probabilities are aligned with the environmental switching process. We evaluated the sign of *λ*″(0) in equation 26 across the space of transmission probabilities. When it is positive, sufficiently small mutations that move the phenotypic value of the rare inherited state in either direction away from 0 increase long-term growth.

When selection is only slightly stronger than *η*^2^ = 1*/*2, the threshold for randomization, phenotypic variability is favored for a restricted range of transmission strategies (Figure 3). For short environmental runs, the ancestral inherited state must be retained with high probability, whereas the alternative inherited state must have a low retention probability. The population therefore continues to produce the ancestral intermediate phenotype most of the time while only occasionally producing the new phenotypic variant.

**Figure 3.**
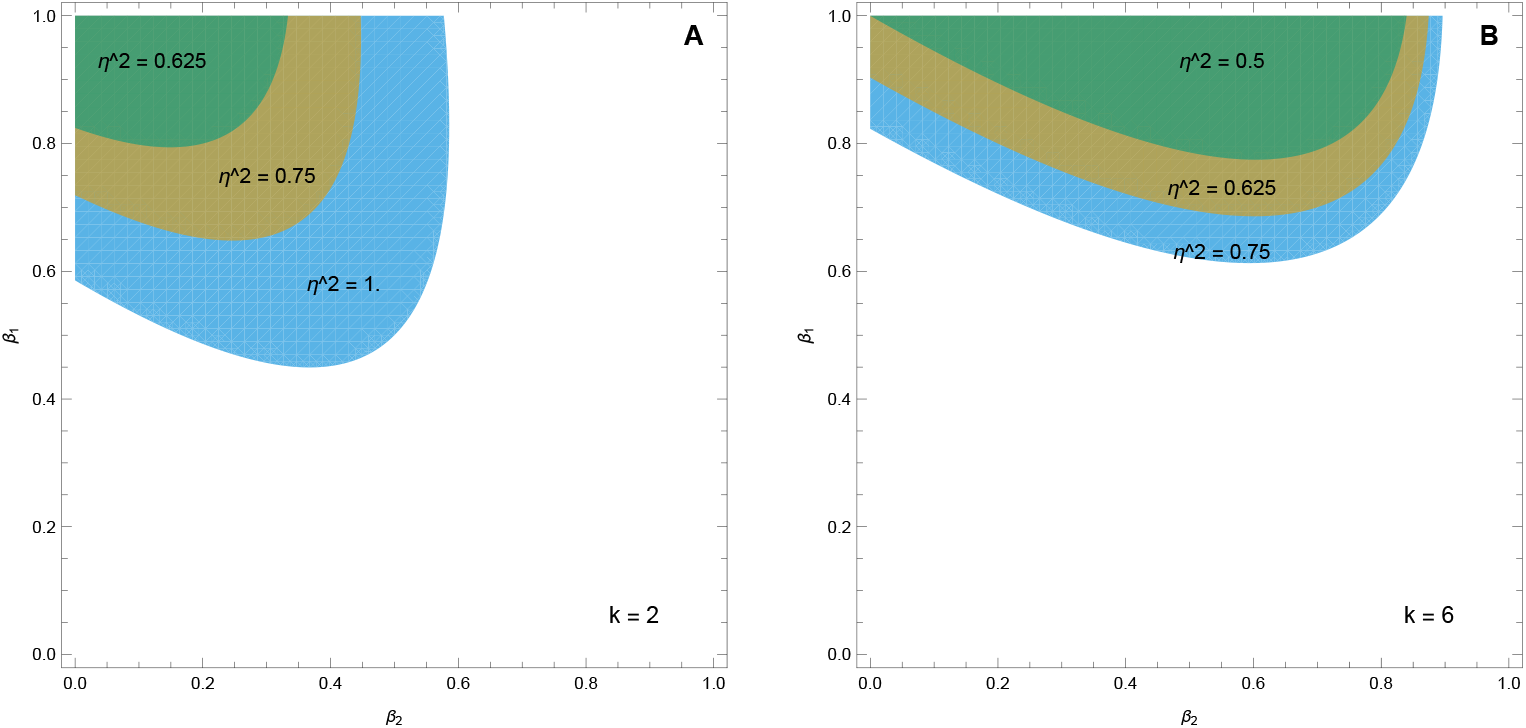
Retention probabilities that permit weak selection for phenotypic variability. Shaded regions indicate combinations of *β*_1_ and *β*_2_ for which the second derivative of invasion fitness is positive. Panel A shows the result for *k* = 2 while panel B shows the result for *k* = 6. The range of transmission strategies permitting invasion increases with *η*^2^ and with the duration *k* of each environmental state.

Stronger selection against mismatch broadens the region of the space of transmission probabilities in which variability can invade. Longer environmental runs also broaden the permissible range of retention for the alternative inherited state. In all cases, however, the initial favorable strategies remain asymmetric, with one inherited state common and reliably transmitted while the second is initially rare and has more restricted transmission. The transmission strategies that permit the initial evolution of phenotypic variability to spread are not the symmetric strategies with high retention of both inherited states, which are, as we show next, favored after the phenotypes have become more different. Expressed in terms of stationary frequency and lag-one correlation, these strategies require the ancestral inherited state to be common, while *h* can be weakly positive or weakly negative for short runs and must generally be positive for longer runs (Supplementary material, section S1 and figure S1). The magnitude of second-order selection also varies within the favorable region and is greatest along a narrow ridge on which the rare inherited state is retained with intermediate probability (Supplementary material, section S2).

### Retention probabilities matched to environmental duration

The preceding results describe invasion near the single-phenotype ancestral strategy. We next considered what happens when two phenotypes are already quite diverged and strongly adapted to opposite environments. In the limiting case, *z*_1_ ≈ −1 and *z*_2_ ≈ 1, so

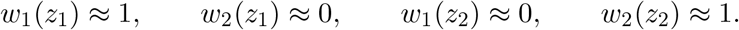

During the *k* − 1 generations following the first generation of an environmental run, growth depends on retaining the currently favored phenotype. In the first generation after an environmental transition, growth instead depends on producing the alternative phenotype. The per-generation log growth rate is therefore

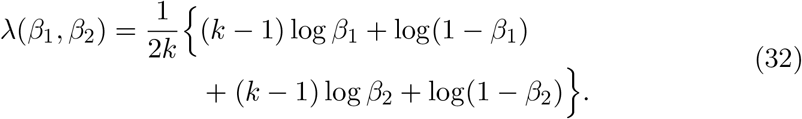

Because the terms involving *β*_1_ and *β*_2_ are independent and symmetric, growth is maximized for

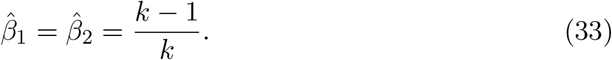

The corresponding probability of switching inherited state is 1*/k*, which matches the frequency of environmental transitions. The same value is the optimal mutation rate of a genetic locus in a periodic environment (Ishii et al. 1989) and the optimal rate of spontaneous phenotypic transitions (Lachmann and Jablonka 1996). The corresponding inherited-state autocorrelation is *ĥ* = (*k* − 2)*/k* = *ρ*, equal to the lag-one autocorrelation of the environment. Substituting the optimal transmission probabilities into equation 32 gives

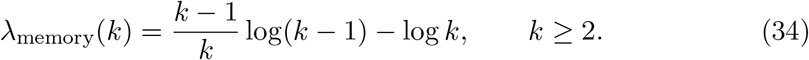

This log growth rate increases with *k* and approaches zero as the duration of each environment becomes long.

The log growth rate of the ancestral single-phenotype strategy is 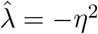 and is strictly less than zero for *η*^2^ *>* 0. Consequently, for any fixed positive strength of selection, the state-transmission strategy eventually exceeds the single-phenotype strategy as *k* increases. For example, when *η*^2^ = 1*/*2, the two strategies have approximately equal log growth at *k* = 5, and the state-transmission strategy has higher log growth for longer environmental runs (Figure 4). This comparison illustrates that a mutant producing highly diverged phenotypes with state transmission can have a large fitness advantage even though selection at its origin is weak.

**Figure 4.**
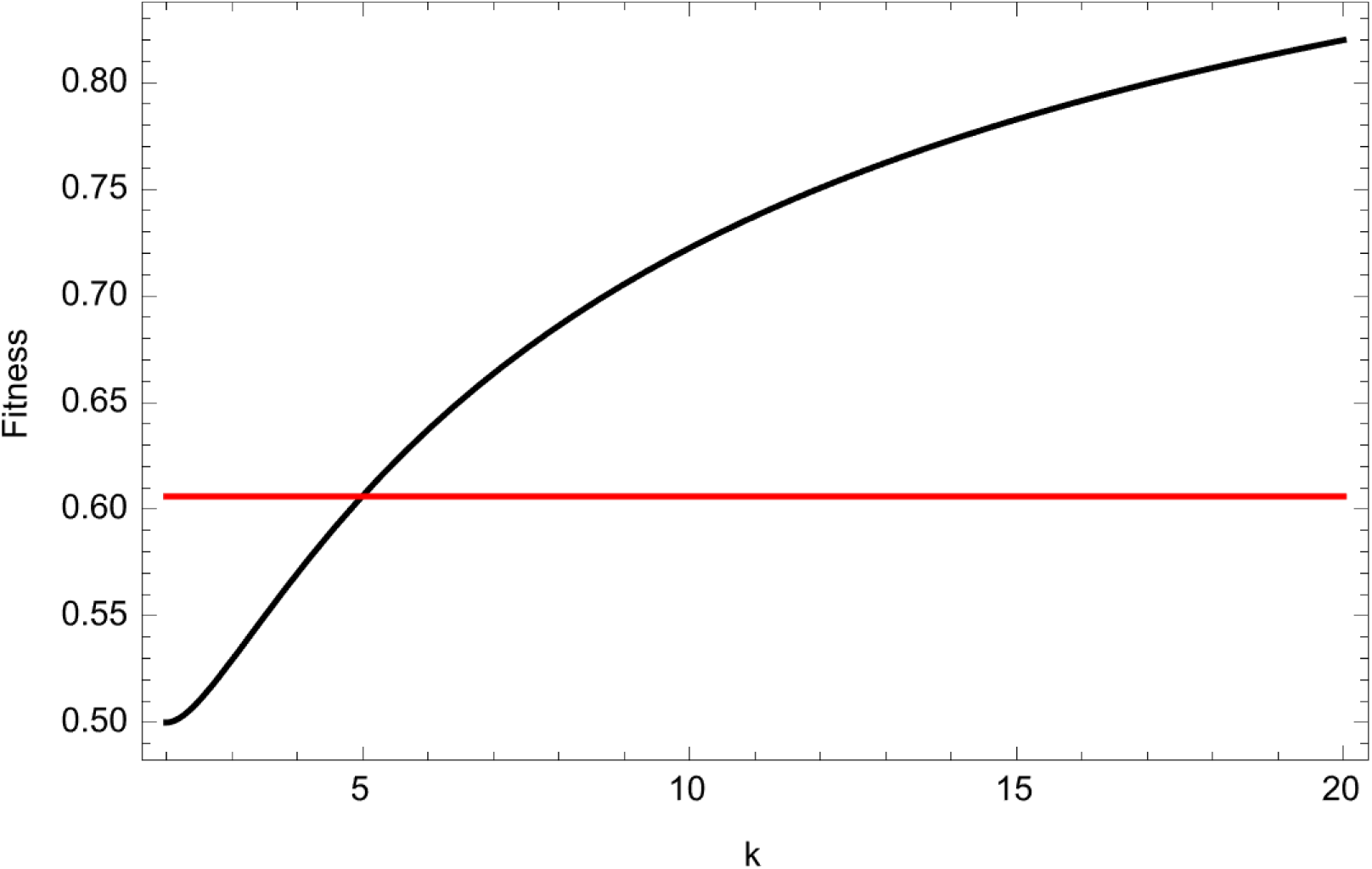
Geometric mean fitness of a state-transmission strategy with two phenotypes strongly adapted to opposite environments. The black curve shows the geometric mean fitness, *e*^*λ*^, when both inherited states have a retention probability of (*k* − 1)*/k* (equation 34). The horizontal line shows the geometric mean fitness of the ancestral single-phenotype strategy when *η*^2^ = 1*/*2.

### Joint evolution of phenotype and transmission

We simulated evolutionary trajectories to examine how the two phenotypic values and their retention probabilities evolve together, and to see how the fitness gradient changes as the amount of phenotypic differentiation increases. The simulations began with one common phenotype at the ancestral equilibrium and one rarely produced phenotype with a small phenotypic difference, so that ***θ*** = (*z*_1_ = 0, *z*_2_ = 0.10, *β*_1_ = 0.99, *β*_2_ = 0.01). Figure 5 shows a characteristic trajectory for *k* = 6 and *η*^2^ = 0.75.

**Figure 5.**
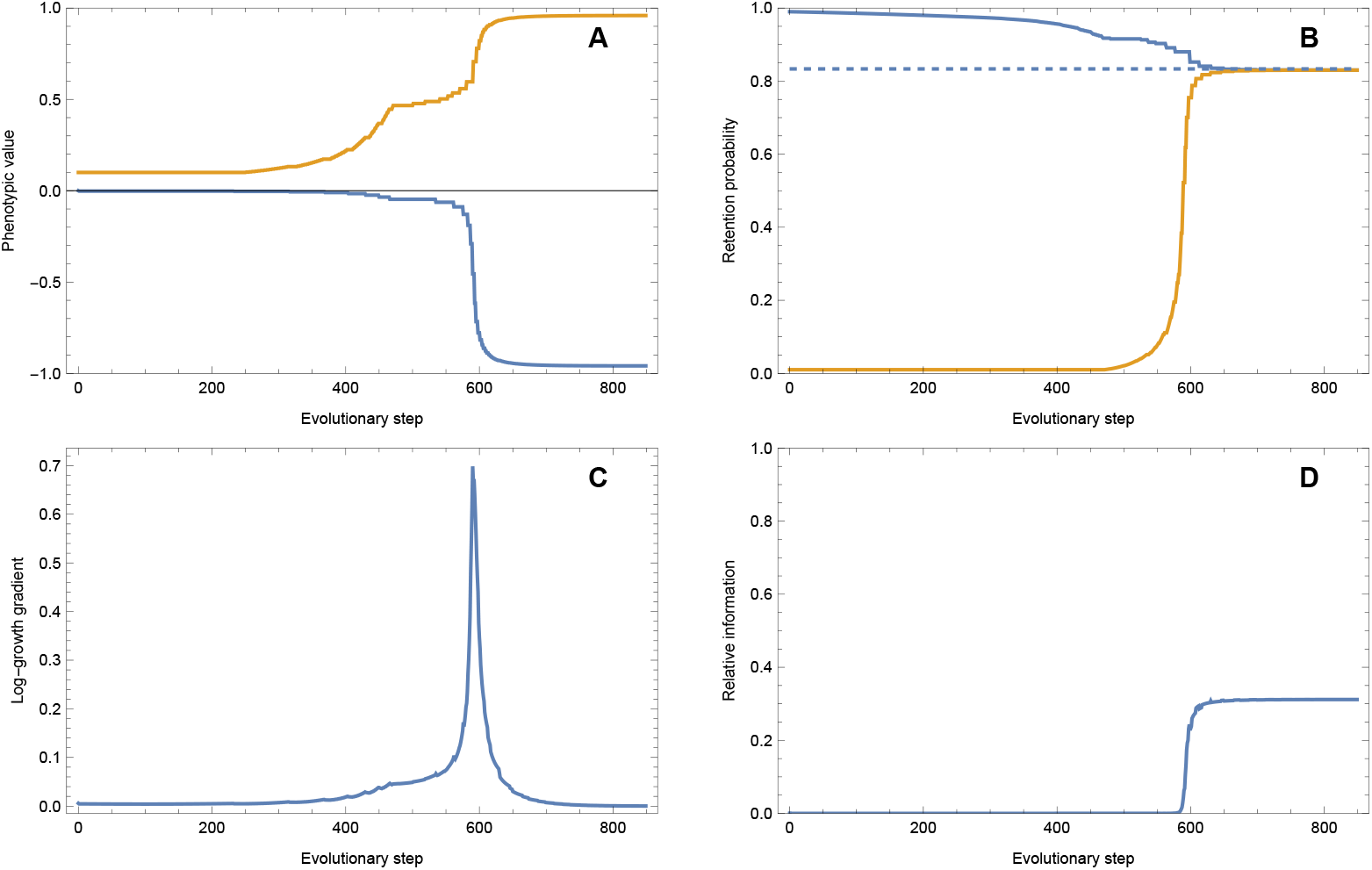
Evolutionary trajectory of phenotype and state transmission. We simulated evolution by calculating the local fitness gradient and changing the component of the strategy with the largest absolute derivative. Parameters are *k* = 6 and *η*^2^ = 0.75, with initial conditions *z*_1_ = 0, *z*_2_ = 0.10, *β*_1_ = 0.99, and *β*_2_ = 0.01. Panel A shows the phenotypic values, with *z*_1_ in blue and *z*_2_ in orange. Panel B shows the retention probabilities, with *β*_1_ in blue and *β*_2_ in orange. The dashed line marks (*k* − 1)*/k*, the optimal retention probability for strongly differentiated phenotypes. Panel C shows the magnitude *G* of the fitness gradient, and Panel D shows mutual information between inherited state and environment relative to the maximum attainable under state transmission.

The fitness gradient is initially weak, as predicted by the analytical calculations. The trajectory then proceeds through three qualitative phases. In the first, *z*_2_ begins to increase while *β*_1_ decreases, increasing production of the alternative inherited state. During the second phase, after *z*_2_ has diverged substantially from 0, changes in *β*_2_ begin to have appreciable fitness effects, and the two retention probabilities evolve in response to one another.

In the final phase, *z*_1_ begins to evolve to levels below 0 while *z*_2_ moves rapidly toward one. The two phenotypes therefore become increasingly specialized on opposite environments. At the same time, *β*_1_ and *β*_2_ converge toward (*k* − 1)*/k*, the optimum for highly differentiated phenotypes. This phase is characterized by a rapid increase in the magnitude of the fitness gradient. Selection becomes stronger only after the phenotypes produced by the two inherited states differ by a large amount. The largest gradient encountered along a trajectory increases with *k* and with *η*^2^ (Supplementary material, section S3).

Relative mutual information between inherited state and environment also increases during this final phase. This increase indicates that the distribution of inherited states becomes more strongly associated with the environmental cycle (see also Chatterjee et al. 2025). Because mutual information depends on the association between inherited state and environment rather than on the quantitative difference between *z*_1_ and *z*_2_, it rises as transmission begins to reliably place the differentiated inherited states in the environments where they are favored. Mutual information is shown relative to the maximum attainable given that the change in environment cannot be detected before it occurs (leading to a one-generation lag).

### The conditions for the maintenance of state transmission

Our results show that the initial spread of small phenotypic differences requires *η*^2^ *>* 1*/*2 for phenotypic randomization. Once diverged phenotypes and state transmission have evolved, however, they may remain favored under weaker selection. We examined this possibility by following the evolved strategy while changing *η*^2^ in small steps up and down.

To compare the conditions for origin and maintenance, we changed *η*^2^ over a range from 0.125 to 1. We scanned up from weak selection and increased *η*^2^, and also scanned down, starting with *η*^2^ = 1 and then decreasing it. At each value of *η*^2^, the simulation was initialized with the evolved strategy obtained at the preceding value and the simulation continued until the largest absolute component of the gradient was less than 10^−4^. When a simulation ended with effectively no phenotypic variation, a small amount of variation was reintroduced before continuing the scan so that the trajectory did not remain fixed at an exactly monomorphic strategy (Supplementary material, section S4).

The resulting equilibria show clear hysteresis, in that the equilibrium reached depends on the direction from which *η*^2^ is changed (Figure 6). Beginning from the ancestral single-phenotype strategy, differentiated phenotypes and state transmission evolved only after selection became sufficiently strong. Beginning instead with established state transmission, differentiated phenotypes and their transmission persisted as selection was weakened below the conditions that permitted their origin.

**Figure 6.**
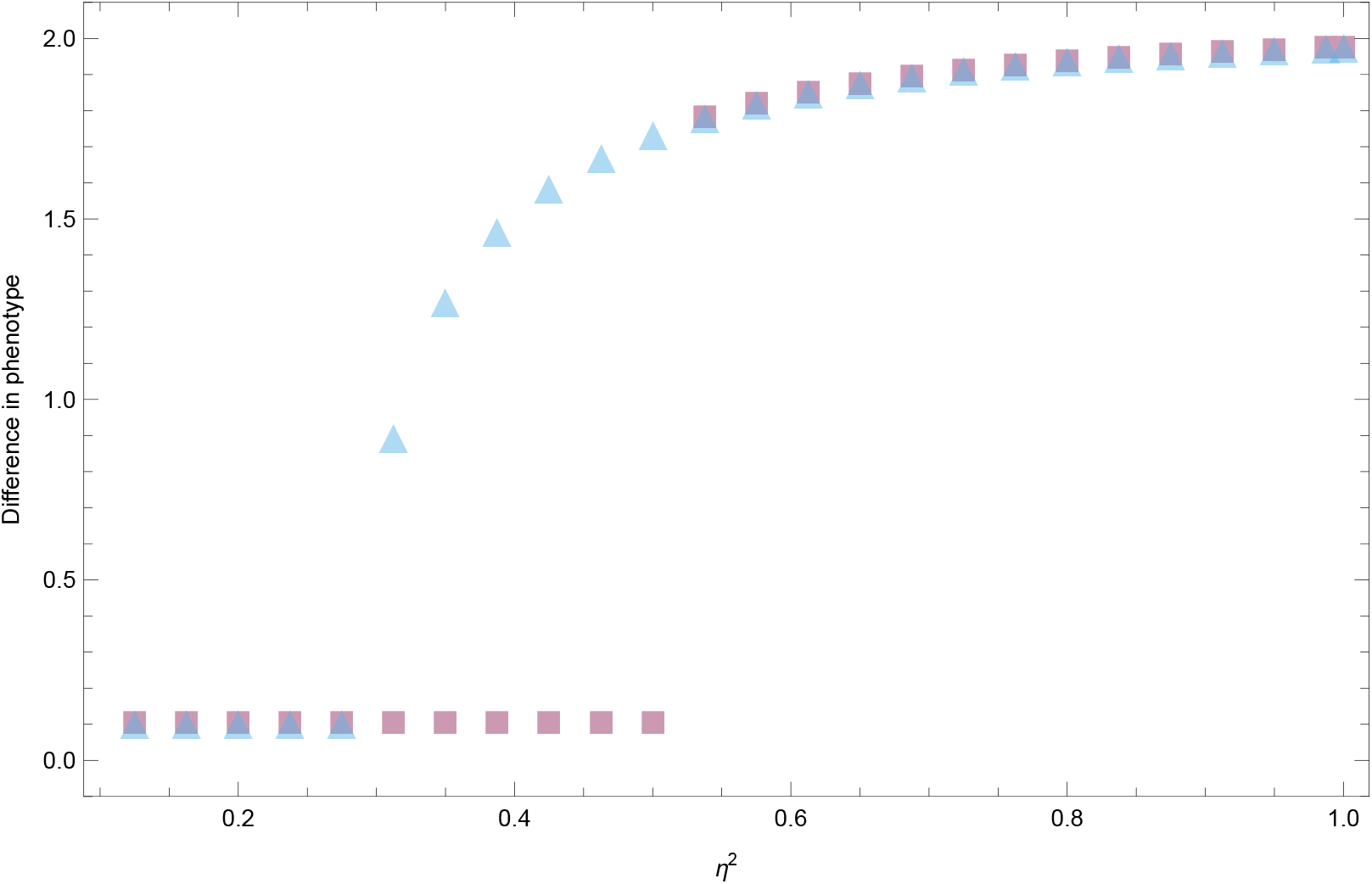
Hysteresis in evolved phenotype and state transmission as the strength of selection against phenotype–environment mismatch changes. Environmental runs last *k* = 6 generations. Squares show the scan that begins at *η*^2^ = 0.125 and proceeds toward *η*^2^ = 1. Triangles show the scan that begins at *η*^2^ = 1 and proceeds in the opposite direction. Each simulation begins from the equilibrium reached at the preceding selection strength. The vertical axis is the difference between the two evolved phenotypic values, *z*_2_ − *z*_1_.

## Discussion

Our main result is that in periodically fluctuating environments, the transmission of inherited states, and the phenotypic memory it generates, cannot evolve by small steps from a phenotypically monomorphic population. At the ancestral equilibrium a small difference between the phenotypic values of the two inherited states has no first-order effect on long-term growth, regardless of the retention probabilities, because these have no fitness effect until the inherited states produce different phenotypic values. Whether the difference spreads depends on second-order effects, and it spreads only when selection against phenotype–environment mismatch is strong (Figures 1 and 3). The strategies that can then invade are asymmetric, with one inherited state common and close to the ancestral phenotypic value and the other rare and, for short environmental runs, poorly retained (Figure 3). Selection strengthens only as the two phenotypic values diverge, and genotypes with retention probabilities matched to the duration of the environments are then favored (Figures 4 and 5). Once differentiated phenotypic values and their transmission have evolved, they persist when selection is weakened below the strength required for their origin from a randomizing ancestor (Figure 6). A maternal effect conditioned on the maternal environment is, in contrast, under first-order selection from the start (Figure 2).

The transmission matrix we analyzed is a general description of two-state inheritance. Read as a model of inheritance, it is a mutation matrix with evolvable and asymmetric rates 1 − *β*_1_ and 1 − *β*_2_, and the retention probabilities are modifiers in the sense of Karlin and McGregor (1974) and Feldman and Liberman (1986), loci that change how other variation is transmitted without affecting fitness on their own, and they are favored only through the associations they build between inherited states and the environment (Liberman et al. 2011; Carja et al. 2014). Population-genetic models of “epialleles”, heritable states of a locus that differ in a molecular mark rather than in sequence, reach the same conclusion from the other side, in that the fate of an epiallele depends on its rate of reversion and on the genetic background that controls it (Slatkin 2009; Geoghegan and Spencer 2013; Furrow and Feldman 2014; Webster and Phillips 2024). The description does not depend on the molecular nature of the inherited state. It applies to physiological conditions, molecular marks and genetic loci with high mutation rates alike, because the transmission of an inherited state is an inheritance system with its own fidelity that leaves the rules of selection unchanged (Haig 2007; Shea et al. 2011; Charlesworth et al. 2017). The term “epigenetic”, whose meanings have multiplied (Deans and Maggert 2015), plays no part in the selection argument.

Because our model is a first-order Markov process (Methods), the lag structure of a time series follows from the one-generation transmission. If the inherited state has components that are not part of the measured phenotype, the same one-generation process gives a different lag structure, in which correlations can alternate in sign across generations or be larger at lag two than at lag one, as the multivariate maternal-effect models of Kuijper et al. (2014) show for continuous phenotypes. An exposure three generations earlier can therefore affect offspring through the inherited states transmitted across the intervening generations without being a separate input to development. A record that distinguished particular ancestral generations would itself have to be carried in the inherited state received by the offspring.

Reports that an inherited state produced in response to the environment is expressed for a fixed number of generations (Heard and Martienssen 2014; Klosin et al. 2017; Bošković and Rando 2018; Houri-Zeevi et al. 2020) are therefore not evidence for a process that records the number of generations. The same description applies to the delayed impact of stress, “transgenerational” plasticity and “priming”, non-genetic responses that unfold over several generations and are therefore difficult to distinguish from the response to selection on genetic variants (Zilio et al. 2026), and to the carry-over within lineages that changes population growth and extinction risk in fluctuating environments (Rescan et al. 2020; Abreu et al. 2024). All of these are one-generation processes with a retention probability, so that the stability and reversibility across generations that Zilio et al. (2026) propose to measure are the retention probabilities themselves. Because the retention probabilities are equivalent to the stationary frequency of an inherited state and its lag-one correlation (equations 7 and 8), a phenotypic time series is enough to estimate them. We did so from population growth in *C. elegans* under fluctuating oxygen (Proulx et al. 2019), where the asymmetric retention inferred is of the kind the model predicts at the origin of state transmission (Figure 3). A time series does not, however, identify the process that generates the carry-over. Environment-independent retention, a maternal effect that persists in part into the next generation and plasticity to a lagged cue can all produce the same pair of values. What distinguishes them, the order of selection at their origin, is not observable in such a series.

Our analysis rests on four assumptions. We considered mutations of small effect. A mutation of large effect could bypass the region of weak selection, but only if it produced an appropriate alternative phenotypic value and produced it at an appropriate frequency. We further considered an asexual population, in which parent and offspring have identical fitness interests, so that the question of whose phenotype a retention probability is does not arise (Wolf and Wade 2001; Hadfield 2012; Wolf and Wade 2016). With sexual reproduction, selection on the mother and selection on her offspring can favor different retention probabilities (Uller and Pen 2011; Kuijper and Johnstone 2018), and the build-up of a correlation between the inherited state of the mother and the environment depends on whether selection acts on viability or on fecundity (Kuijper and Johnstone 2021). We also assumed that transmission was independent of the environment. Models in which the inherited state is reset during development toward the phenotype favored in the current environment (Jablonka et al. 1995; Lachmann and Jablonka 1996; Uller et al. 2015) take as given that the environment is sensed, that the reset goes in the direction the environment favors and that the alternative phenotypic values already exist. A single mutation is unlikely to produce the three at once, and how they arise in sequence from an ancestor that has none of them should be the subject of further work.

The last assumption is that the environment in our model alternates periodically, with runs of *k* generations, so that the sequence of past environments determines the future one. When the environment instead changes with a constant probability each generation, the maternal environment predicts the distribution of offspring environments, and earlier environments add nothing to the prediction. Environments that are mostly periodic lie between the two. The absence of first-order selection at the origin does not depend on periodicity, however, because at the ancestral equilibrium the inherited states are not associated with the environment, regardless of its sequence, and first-order selection on a maternal effect conditioned on the maternal environment depends only on the lag-one autocorrelation of the environment (equation 29). The contrast between the two kinds of mutation therefore holds whether the environment changes periodically or at random. Once the phenotypic values have diverged, random environments are more difficult to analyze, because the association between inherited state and environment that selection builds up is then itself random, and long-term growth depends on the whole sequence of environments rather than on a single cycle.

The contrast between mutations whose phenotypic effect depends on a cue and mutations whose effect does not can also be framed in terms of information. Retention probabilities do not depend on the environment, so a lineage without cues acquires information about the environment only through differential survival or reproduction of the inherited states it carries (Kuijper and Johnstone 2021). What state transmission preserves is the result of past selection. Mutual information between inherited state and environment (see Methods) describes this association, but selection does not maximize it, because the same amount of information can go with phenotypic effects of any direction and size (Donaldson-Matasci et al. 2010; Rivoire and Leibler 2011; Dall et al. 2015; English et al. 2015). In our simulations, information and the strength of selection rise together only because the evolving retention probabilities come to place the differentiated phenotypic values in the environments where they are favored (Figure 5). Describing maternal effects as anticipatory (Marshall and Uller 2007; Storm and Lima 2010) refers to the same correlation, between the cue the mother receives and the selection that offspring go on to experience (Leimar and McNamara 2015; Nettle and Bateson 2015; McNamara et al. 2016).

The contrast between first- and second-order selection also separates our model from the quantitative-genetic description of maternal inheritance. In the maternal-inheritance model of Falconer (1965) and Kirkpatrick and Lande (1989), the maternal phenotype enters the offspring phenotype through a coefficient *m*, the regression of offspring on maternal phenotype that remains once genetic transmission is accounted for, and the lag-one correlation *h* between the inherited states of parent and offspring plays the same role for a two-state phenotype (Table 1). In that model the phenotype already varies among mothers, so a small change in *m* changes the phenotypes of their offspring in proportion to how much each mother deviates from the mean. The fitness effect of the change is first order whenever the maternal phenotype carries information about the offspring environment. The benefit of a given *m* accordingly depends on the predictability of environmental change (Hoyle and Ezard 2012). When *m* is allowed to evolve in a periodically fluctuating environment it evolves according to the autocorrelation between maternal and offspring environments, positive when change is slow and negative when it is rapid (Kuijper and Hoyle 2015). When maternal effects involve several traits, differences in the timing of selection on those traits can also favor cross-trait maternal effects (Kuijper et al. 2014). The retention probabilities matched to the duration of the environments that are favored in our model once the phenotypic values have diverged correspond to *h* = (*k* − 2)*/k* (equation 33), the lag-one autocorrelation *ρ* of the environment, so the two descriptions agree on the endpoint and differ in the starting point. The difference in starting point is that the variation that *m* transmits is present from the outset, whereas here it must first arise.

In conclusion, whether a mutation that changes how the phenotype is determined invades depends on whether its phenotypic effect is conditioned on an informative cue when it first arises. Where it does, selection is first order and small mutational steps can build up to large phenotypic differences. Where it does not, as for the transmission of an inherited state that generates phenotypic memory, the first steps are nearly neutral, and the differences in phenotypic value that give transmission its fitness consequences must come from elsewhere, probably from cue-dependent maternal effects.

## Acknowledgments

This work was partly supported by the Agence Nationale de la Recherche (ANR-17-CE02-0017-01 and ANR-24-CE12-0904 to H.T.) and the National Science Foundation (EF-1137835 to S.R.P.). S.R.P.: conceptualization, formal analysis, software, visualization, writing—original draft, review and editing; H.T.: conceptualization, funding acquisition, writing—review and editing. OpenAI ChatGPT, using GPT-5.6 Sol with the Ultra reasoning setting (last accessed 14 September 2026) was used to check the derivations of the equations for errors and the notation for consistency. Anthropic Claude Fable 5.1 with the Max reasoning setting (last accessed 4 October 2026) was used to provide suggestions for text editing. We reviewed and verified all output from these tools and take responsibility for the content of the article. We declare we have no competing interests.

## Supplementary Results

### S1. Transmission strategies expressed as phenotype frequency and lag-one correlation

The transmission matrix used in the main analysis can be expressed using the stationary frequency of inherited state 1 and the lag-one correlation of the inherited state (main text, equations 7–9). Figure S1 presents the complete four-panel version of the invasion-region analysis of main-text Figure 3. The upper panels show the results in the native *β*_1_–*β*_2_ parameterization, while the lower panels show the same regions in terms of stationary phenotype frequency and lag-one correlation.

For short environmental runs (*k* = 2), selection only slightly stronger than *η*^2^ = 1*/*2, the threshold for randomization, favors a narrow set of strategies in which the ancestral intermediate phenotype is common and the alternative inherited state is produced rarely. On the frequency–correlation scale, the ancestral inherited state must have high frequency, while the lag-one correlation can be weakly positive or weakly negative. Increasing *η*^2^ broadens the invasion region, but the favorable strategies remain asymmetric, in that the ancestral inherited state is common and reliably reproduced whereas the alternative inherited state is initially less persistent. For longer runs (*k* = 6), a broader range of values of *β*_2_ permits the alternative phenotype to begin diverging. Under selection only slightly above the threshold, *β*_1_ must remain high, but *β*_2_ can vary substantially. The frequency–correlation representation shows that these strategies generally require positive phenotypic correlation unless the ancestral inherited state is extremely common. Stronger selection again expands the favorable region. Thus, longer environmental runs make inherited persistence more useful, but do not remove the asymmetry characteristic of the origin of phenotypic variation.

### S2. The magnitude of selection varies across transmission strategies

The invasion conditions show when the fitness of a mutant, as approximated by a Taylor’s series, is positive. Here we plot the second derivative of mutant log growth with respect to the phenotypic value of the initially rare inherited state, *λ*″(0) in equation 26, as a measure of the magnitude of that fitness difference. The surface in Figure S2 shows that the favorable region contains a narrower ridge on which the magnitude of second-order selection is greatest.

Along this ridge, retention of the common ancestral inherited state can remain high, whereas retention of the rare alternative inherited state is more constrained. The largest fitness effect occurs near *β*_2_ ≃ 0.8, close to the retention probability predicted for already differentiated phenotypes when (*k* − 1)*/k* = 0.833. Increasing *β*_2_ substantially beyond this ridge causes the local advantage of the new variant to fall rapidly.

### S3. Maximum selection encountered along evolutionary trajectories

The main text illustrates one evolutionary trajectory, but the amount of selection generated after phenotypic differentiation depends on both the strength of selection against mismatch and the duration of each environmental state. We therefore simulated trajectories across a range of *k* and recorded the largest magnitude of the four-dimensional log-growth gradient encountered before convergence:

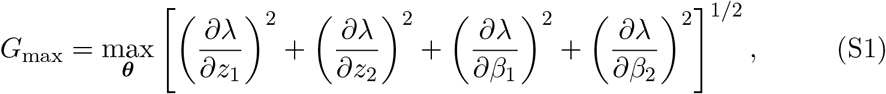

where the maximum is taken over the strategies visited along the trajectory.

**Figure S1.**
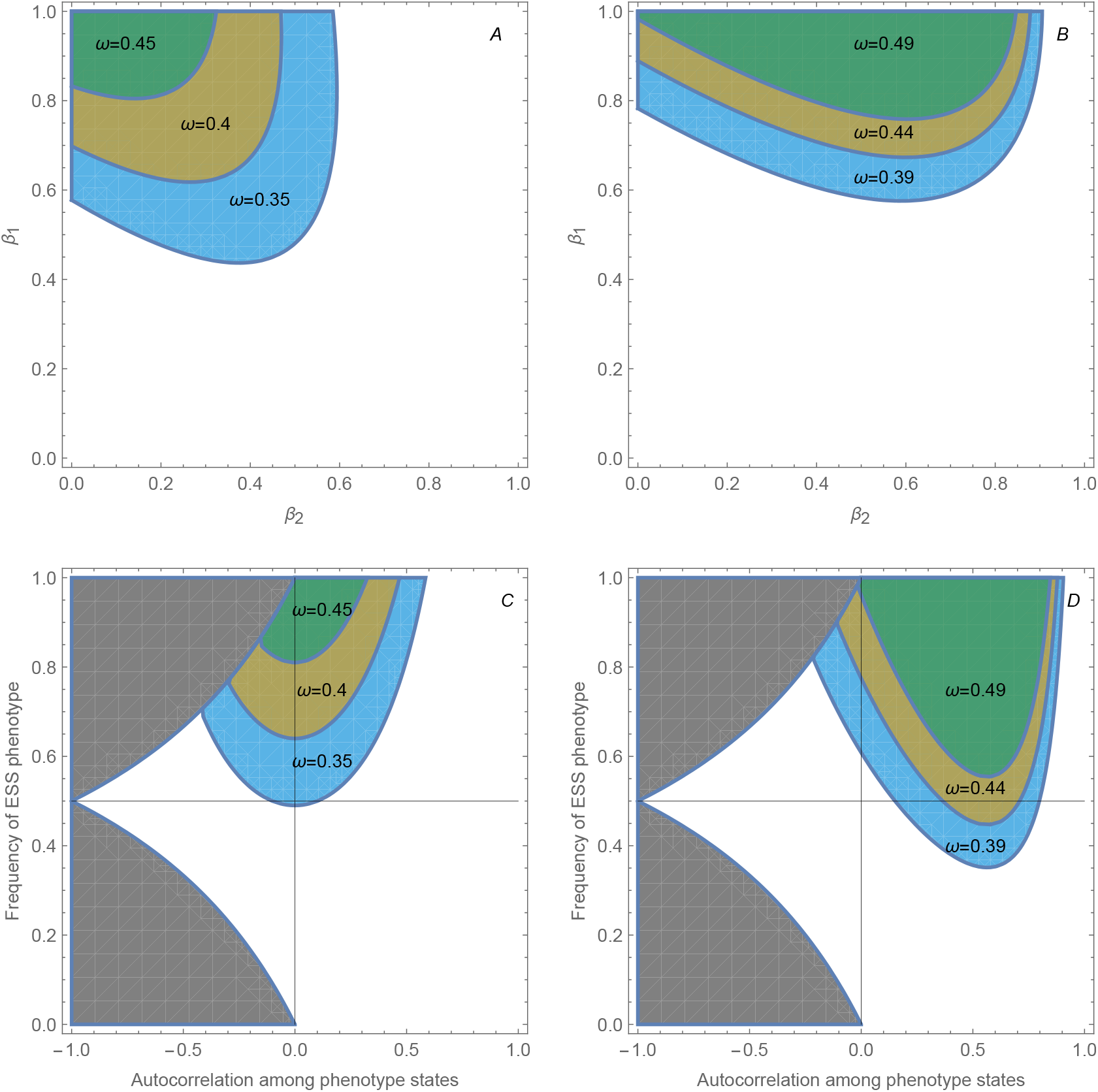
Alternative representations of the transmission strategies that permit weak selection for phenotypic variability. Panels A and B show combinations of *β*_1_ and *β*_2_ for which the relevant second derivative of invasion fitness is positive. Panels C and D show the same regions in terms of the stationary frequency of the ancestral inherited state and the lag-one correlation *h* between the inherited states of parent and offspring. The grey area is the region of the (*h, q*_1_) plane that no transmission matrix can produce. Panels A and C use *k* = 2 and panels B and D use *k* = 6. Curves correspond to different strengths of selection against phenotype–environment mismatch, labelled by *η*^2^.

**Figure S2.**
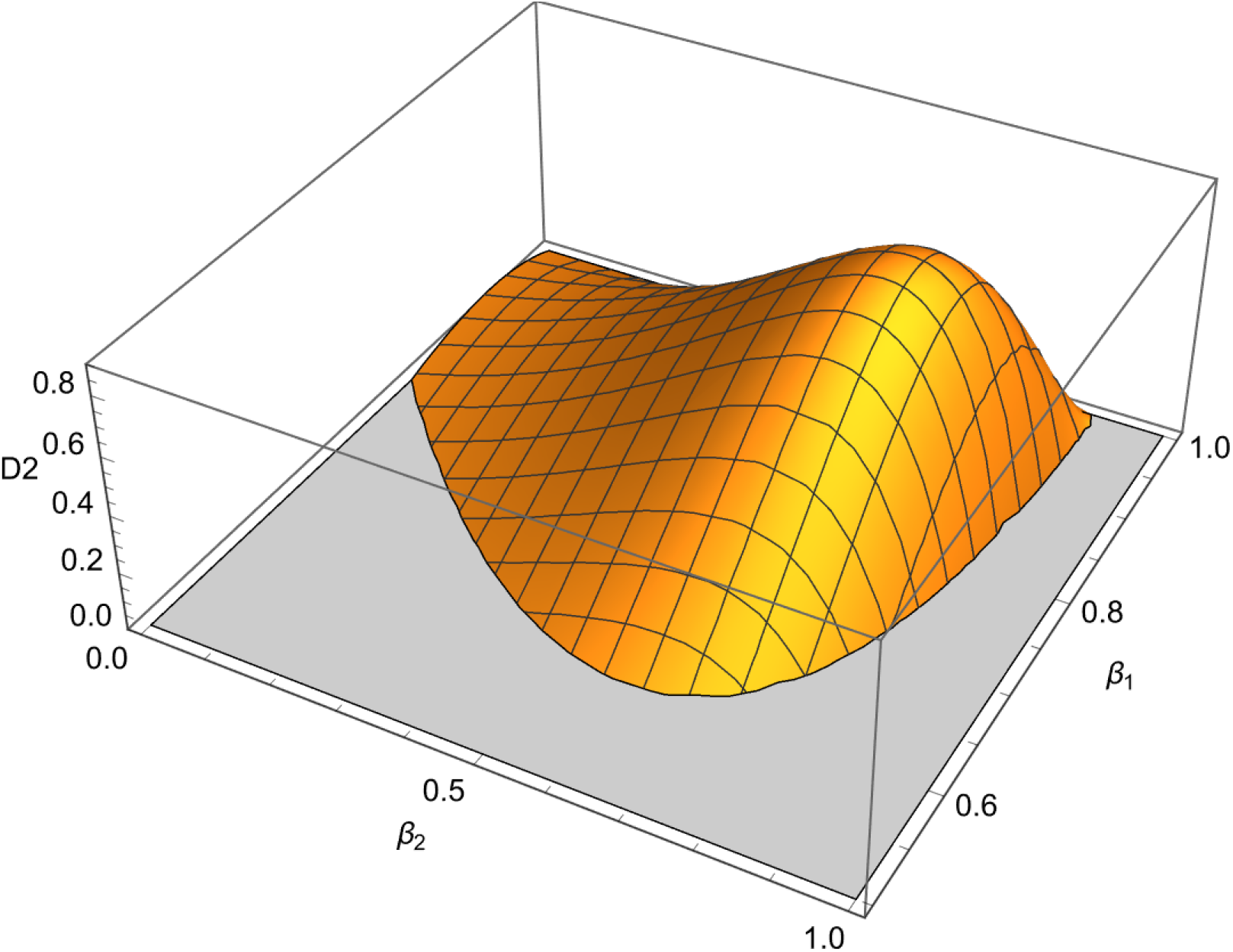
Magnitude of second-order selection across the space of transmission probabilities. The surface shows the second derivative of mutant log growth with respect to the phenotypic value of the initially rare inherited state, *λ*″(0) in equation 26, restricted to the region in which it is positive. Parameter values: *k* = 6, *η*^2^ = 1.

Figure S3 shows that the strongest selection encountered during the evolutionary transition is not determined by the initial derivative, which is zero, but by the strategy reached after phenotype and transmission begin to coevolve. The dependence of *G*_max_ on *k* also emphasizes that a longer environmental run can amplify the consequences of retaining a differentiated phenotype. Stronger mismatch selection produces a larger peak gradient because divergence of the two phenotypes creates larger fitness differences among transmission strategies.

### S4. Additional details of the origin–maintenance comparison

The hysteresis analysis used the equilibrium reached at one selection strength as the initial condition for the next value. For *k* = 6, one scan began at strong mismatch selection and progressively weakened selection, whereas the second began under weak selection and progressively increased it. This continuation procedure follows a local evolutionary branch and is therefore intentionally different from independently restarting the simulation from the ancestral strategy at every value of *η*^2^.

**Figure S3.**
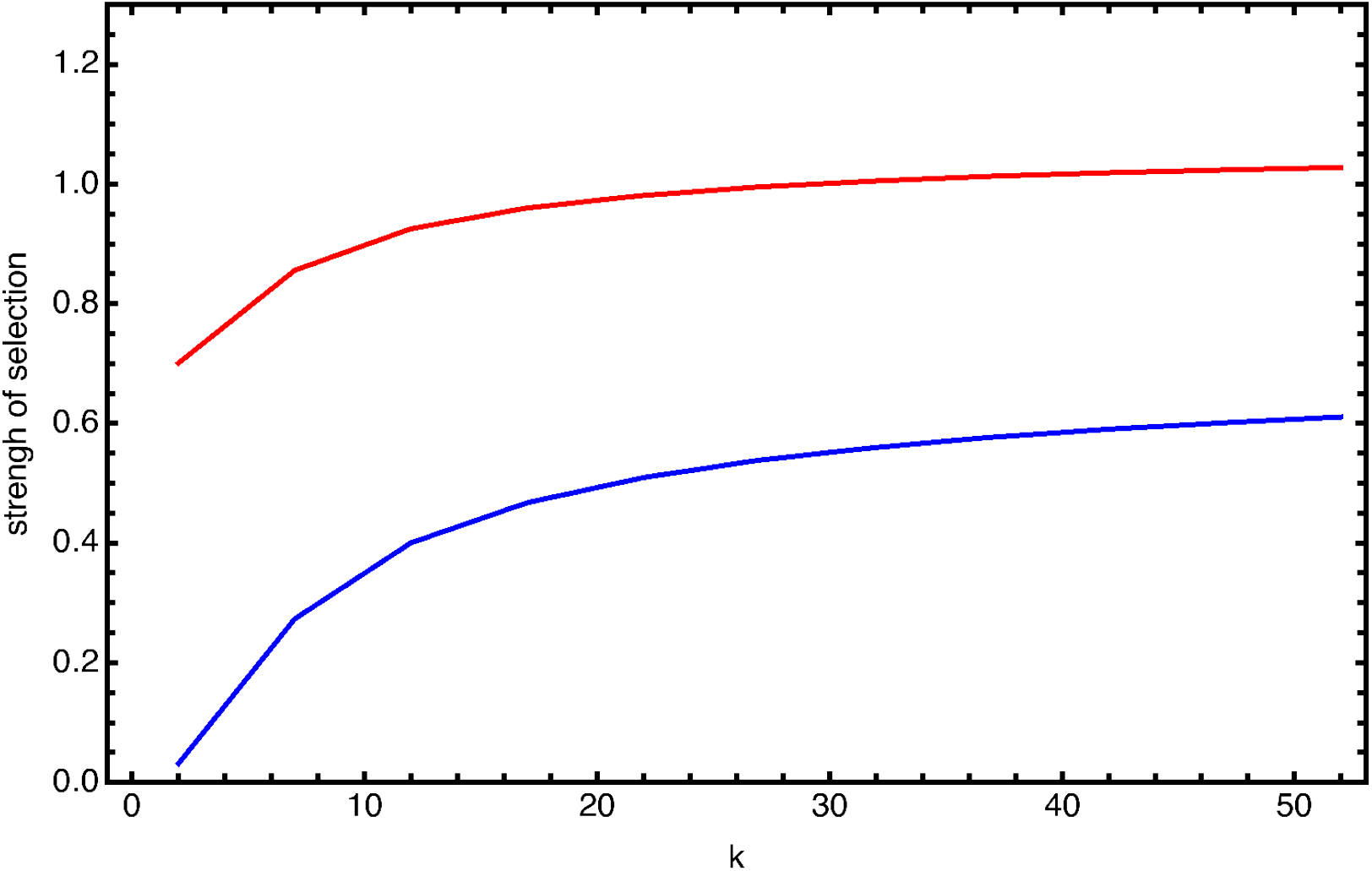
Maximum magnitude of the fitness gradient encountered during evolutionary trajectories as a function of environmental run length *k*. Each trajectory begins near the ancestral single-phenotype strategy and is continued until the local gradient falls below the convergence threshold. The two curves correspond to *η*^2^ ≃ 0.617 (blue; *σ* = 0.90) and *η*^2^ = 2 (red; *σ* = 0.50).

An exactly monomorphic strategy creates a numerical and biological boundary, because once *z*_1_ = *z*_2_ = 0 the transmission probabilities have no effect and a deterministic gradient algorithm cannot regenerate phenotypic variation. During the weak-to-strong scan, we therefore reintroduced the same small, rare phenotypic variant used to initiate the evolutionary trajectories whenever the two phenotypic values were within 0.10 of 0. The reset strategy was

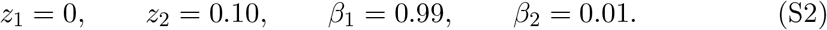

The reset represents a continuing supply of small mutational variation and does not force the variant to spread. The variant disappears again whenever local selection does not support phenotypic differentiation. In contrast, the strong-to-weak scan retains the previously evolved differentiated strategy and asks how far it persists as selection is relaxed. The separation of the two branches therefore reflects the difference between selection acting on a rare, small-effect variant and selection acting on an established state-transmission strategy.

